# Extracellular Vesicle-Mediated Delivery of Genetic Material for Transformation and CRISPR/Cas9-based Gene Editing in *Pneumocystis murina*

**DOI:** 10.1101/2025.06.17.660080

**Authors:** Steven G. Sayson, Alan Ashbaugh, Lillian C. Bauer, George Smulian

## Abstract

*Pneumocystis* species are obligate fungal pathogens that cause severe pneumonia, particularly in immunocompromised individuals. The absence of robust genetic manipulation tools has impeded our mechanistic understanding of *Pneumocystis* biology and the development of novel therapeutic strategies. Herein, we describe a novel method for the stable transformation and CRISPR/Cas9-mediated genetic editing of *Pneumocystis murina* utilizing extracellular vesicles (EVs) as a delivery vehicle. Building upon our prior investigations demonstrating EV-mediated delivery of exogenous material to *Pneumocystis*, we engineered mouse lung EVs to deliver plasmid DNA encoding reporter genes and CRISPR/Cas9 components. Our initial findings demonstrated successful *in vitro* transformation and subsequent expression of *mNeonGreen* and *Dhps*^*ARS*^ in *P. murina* organisms. Subsequently, we established stable *in vivo* expression of *mNeonGreen* in mice infected with transformed *P. murina* for a duration of up to 5 weeks. Furthermore, we designed and validated a CRISPR/Cas9 system targeting the *P. murina Dhps* gene, confirming its *in vitro* cleavage efficiency. Ultimately, we achieved successful *in vivo* CRISPR/Cas9-mediated homologous recombination, precisely introducing a *Dhps*^*ARS*^ mutation into the *P. murina* genome, which was confirmed by Sanger sequencing across all tested animals. Here, we establish a foundational methodology for genetic manipulation in *Pneumocystis*, thereby opening avenues for functional genomics, drug target validation, and the generation of genetically modified strains for advanced research and potential therapeutic applications.

**IMPORTANCE:** *Pneumocystis* species are obligate fungal pathogens and major causes of pneumonia in immunocompromised individuals. However, their strict dependence on the mammalian lung environment has precluded the development of genetic manipulation systems, limiting our ability to interrogate gene function, study antifungal resistance mechanisms, or validate therapeutic targets. Here, we report the first successful approach for stable transformation and CRISPR/Cas9-based genome editing of *Pneumocystis murina*, achieved through *in vivo* delivery of engineered extracellular vesicles (EVs) containing plasmid DNA and encoding CRISPR/Cas9 components. We demonstrate sustained transgene expression and precise modification of the *dhps* locus via homology-directed repair. This modular, scalable platform overcomes a long-standing barrier in the field and establishes a foundation for functional genomics in *Pneumocystis* and other obligate, host-adapted microbes.

## INTRODUCTION

*Pneumocystis* pneumonia (PjP) remains a critical cause of morbidity and mortality in immunocompromised individuals, including those with HIV/AIDS, hematologic malignancies, or organ transplants receiving immunosuppressive therapy ^1-4^. This disease is caused by the obligate fungal pathogen *Pneumocystis jirovecii*. Despite significant clinical relevance, critical aspects of *Pneumocystis* biology, pathogenesis, and antifolate drug resistance remain poorly characterized, primarily due to the persistent inability to culture these fungi *in vitro* ^5^. Unlike other fungi that readily grow in defined media, *Pneumocystis* species are obligate biotrophs, requiring a mammalian host to replicate ^6^, which severely limits functional genetic studies and the exploration of novel therapeutic targets.

A significant clinical challenge associated with *Pneumocystis* treatment is resistance to the antifolate drug combination trimethoprim-sulfamethoxazole (TMP/SMX), which remains the first-line therapeutic option ^7,8^. Mutations in the dihydropteroate synthase (*Dhps*) gene, notably at codons 55 and 57, confer SMX resistance, complicating therapeutic management and necessitating novel methods to dissect resistance mechanisms and validate alternative therapeutic targets ^9-13^.

Furthermore, *Pneumocystis* genomes feature numerous Major Surface Glycoprotein (Msg) genes arranged in tandem repeats near telomeric regions ^14-16^. These Msg genes are hypothesized to undergo frequent genetic recombination, enabling antigenic variation through differential expression of individual Msg variants ^17,18^ The extensive and rapid genetic rearrangements observed in *Pneumocystis* imply a highly efficient DNA recombination system, suggesting that targeted genome editing via homologous recombination may be inherently feasible.

Recently, extracellular vesicles (EVs) have emerged as critical mediators of intercellular communication, transferring proteins, lipids, and nucleic acids between cells ^19^. Our prior work demonstrated that *Pneumocystis* organisms actively internalize host-derived EVs, presumably as a nutrient acquisition mechanism ^20^. We reasoned that this natural EV uptake pathway could be exploited to deliver genetic constructs, facilitating genetic manipulation of *Pneumocystis in vivo*.

CRISPR/Cas9-mediated genome editing technology has transformed genetic engineering, allowing targeted disruption, insertion, or modification of specific genomic loci ^21,22^. Adapting this technology for *Pneumocystis* would significantly enhance our ability to probe gene function, dissect resistance mechanisms, and identify novel therapeutic targets.

In the present investigation, we aimed to develop a comprehensive methodology for genetic manipulation in *P. murina*, encompassing two distinct, yet complementary approaches: stable plasmid-based transformation for gene overexpression and precise CRISPR/Cas9-mediated gene editing for targeted genomic modifications. We hypothesized that engineered mouse lung EVs could effectively deliver plasmid constructs to *P. murina*, enabling stable expression of foreign genes and facilitating targeted genomic modifications. Herein, we present compelling data demonstrating successful *in vitro* transformation, the establishment of *in vivo* persistence of transformed organisms, and the achievement of precise CRISPR/Cas9-mediated homologous recombination at the *Dhps* locus, a critical gene implicated in antifolate drug resistance. This work represents a significant leap forward in *Pneumocystis* research, providing essential tools that will facilitate future functional genomic studies, enable the creation of genetically modified strains, and ultimately contribute to the development of more effective therapeutic strategies against PjP.

## RESULTS

### EV Uptake by *P. murina*

Our previous work demonstrated that *P. carinii* actively uptakes native host lung EVs labeled with the lipophilic dye PKH26, suggesting a mechanism for nutrient acquisition from the host ^20^. Building on this, we conducted an initial feasibility study to determine if *P. murina* could internalize EVs containing exogenous nucleotides. EVs transformed with TxRed-labeled siRNA were co-cultured with *P. murina* for 24 hours.

Fluorescent microscopy analysis (Figure 1) demonstrated clear uptake of the TxRed signal by *P. murina* organisms. In the control group, where *P. murina* was co-cultured with siRNA-TxRed but no EVs, no TxRed signal was observed within the organisms. Similarly, *P. murina* co-cultured with EVs containing no cargo also showed no TxRed signal. In contrast, *P. murina* co-cultured with EVs loaded with siRNA-TxRed exhibited distinct punctate TxRed fluorescence localized within the fungal cells, indicating successful internalization of the EV cargo. This confirms that *P. murina* can actively take up EVs and their contents, establishing a foundational step for EV-mediated genetic delivery.

**Figure 1.**
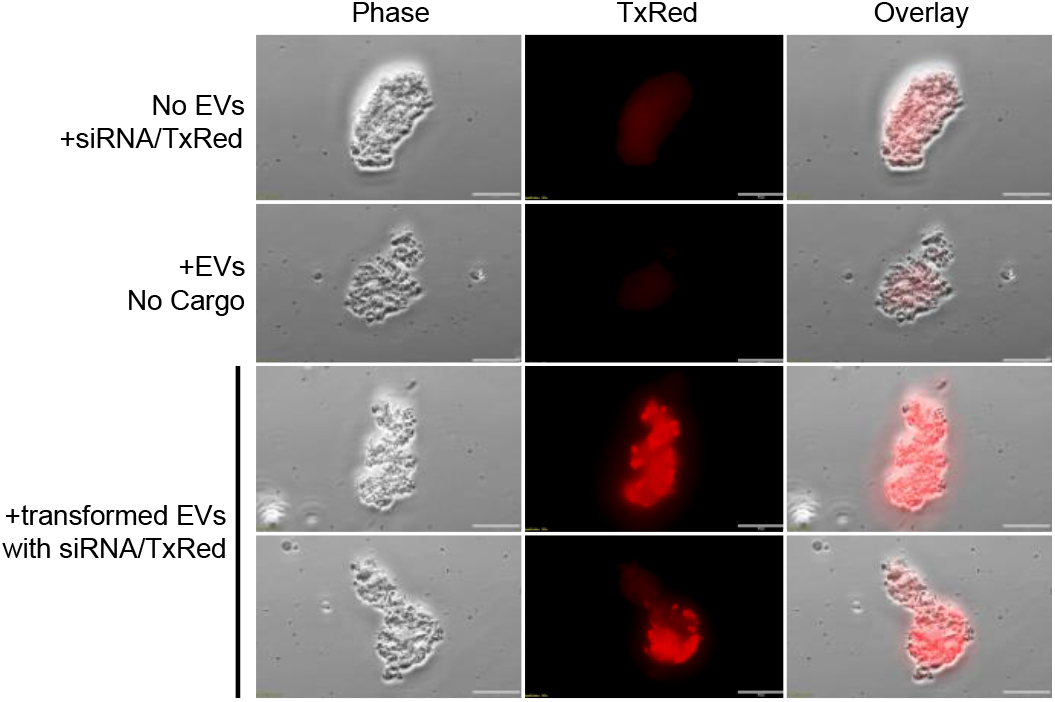
*Pneumocystis murina* uptake BALF EVs containing exogenous nucleotides. *P. murina* were treated with TxRed-conjugated siRNA (top row), EVs alone (middle row), or EVs-loaded with siRNA-TxRed (bottom row) for 16 hours. Scale, 20 μm.

### Successful EV-Mediated Gene Delivery and Expression in *P. murina*

The initial pSS1 plasmid (Figure 2A) was constructed, incorporating an expression cassette with a flanking *P. murina Msg* promoter and *PmNamp8* terminator to ensure the expression of introduced genes within the fungal host. *P. murina* codon optimized *mNeongreen* or *Dhps*^*ARS*^ were cloned into the coding region.

**Figure 2.**
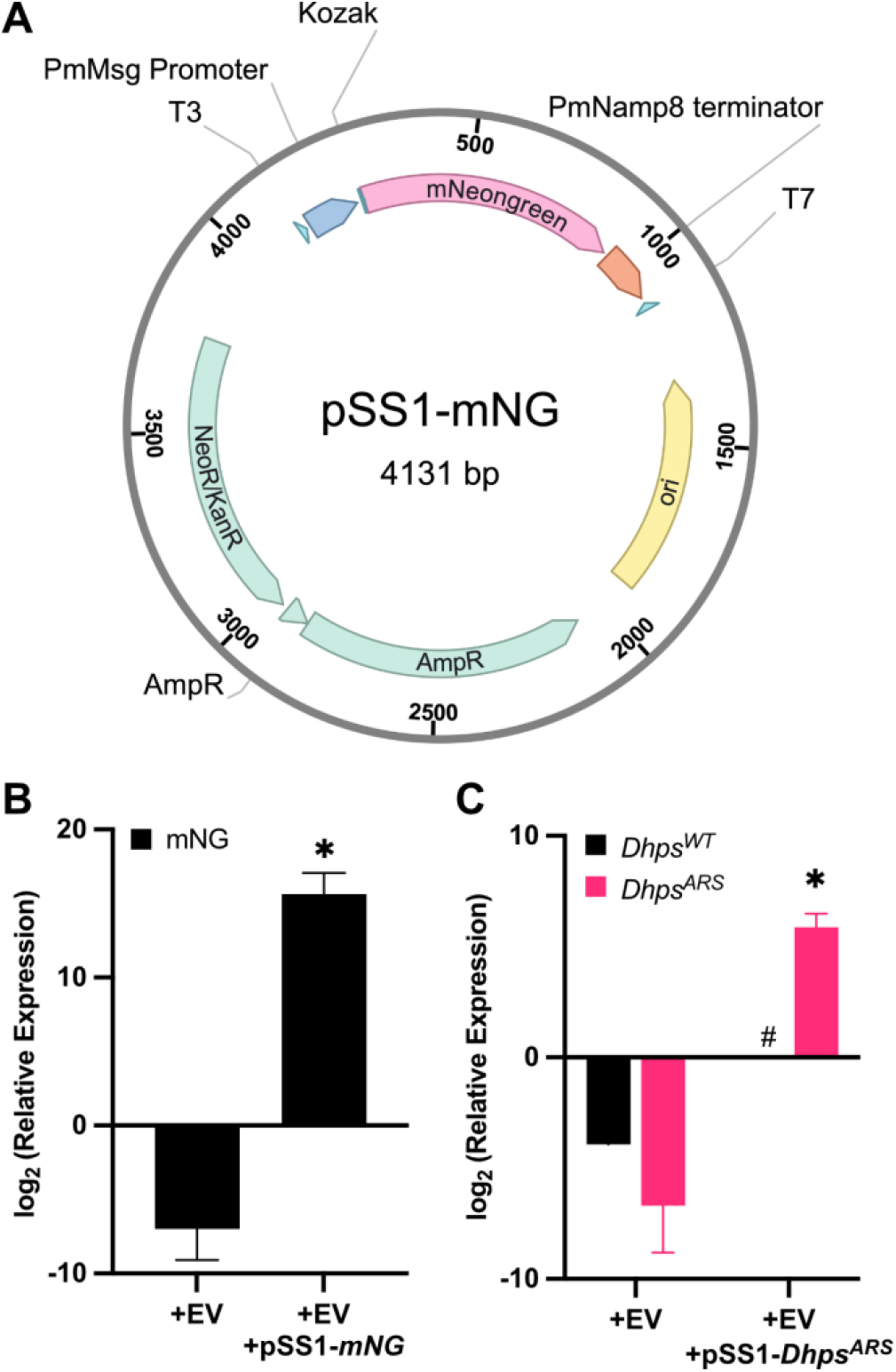
*Pneumocystis murina* cultured *in vitro* transcribes plasmid-encoded genes following delivery by extracellular vesicles. (A) Plasmid map of pSS1-mNG, which encodes mNeongreen driven by the Pm *Msg* promoter, and a Namp8 terminator. P. murina (1 × 10^6^ organisms) were treated with EVs (2-μg protein equivalent) loaded with either pSS1-mNeongreen (mNG) or pSS1-Dhps^ARS^ for 16 hours. Total RNA was extracted from the cells and synthesized to cDNA. RT-qPCR was performed and relative expression was calculated using the 2^-ΔCt^ method, normalized to LSU. Plasmid-transformed (B) mNeongreen and (C) Dhps^ARS^ groups (n=9) showed significant transcriptional activity, confirming successful expression of the plasmid-delivered genes. Native Dhps^WT^ was notably absent in plasmid transformed P. murina. T-test; *, p < 0.05 against control. ^#^, unable to perform comparison due to the lack of detectable expression.

*P. murina* organisms co-cultured with EVs containing either pSS1-*mNeongreen* or pSS1-*Dhps*^*ARS*^ demonstrated successful gene delivery and subsequent transcriptional activity *in vitro* after 24 hours. Relative to *P. murina* exposed to empty EVs (control), the *mNeongreen* and *Dhps*^*ARS*^ groups showed strong transcriptional activity, with average 2^-ΔCt^ values of 51,333 and 58.95, respectively. These correspond to log2-transformed values of approximately 15.7 and 5.9, confirming successful expression of the plasmid-delivered genes (Figure 2B/C). Interestingly, endogenous *Dhps*^*WT*^ transcription was not detected in pSS-Dhps^ARS^ transformed Pm, suggesting a feedback mechanism regulating gene expression. Western blot analysis did not detect mNeongreen protein expression (data not shown), likely due to the lack of a long-term culture system. Under these conditions, *P. murina* is unable to maintain cellular homeostasis, thereby likely limiting its capacity for efficient protein translation.

This initial *in vitro* success established the feasibility of EV-mediated gene delivery for *Pneumocystis* and foundational proof-of-concept for the entire study. It demonstrates that EVs can indeed deliver genetic material to *P. murina* and, crucially, that this material is transcriptionally active within the fungal cells. Without this initial success *in vitro*, the more complex *in vivo* experiments and CRISPR applications would not be feasible.

### Development of a Stable *In vivo* Transformation System

To enable stable and long-term gene expression within the host, the pSS2.1 plasmid (Figure 3A) was engineered. This improved construct incorporates a blasticidin resistance (*bsd*) gene for selective pressure and, critically, a truncated centromeric region, *PmCen15* ^23^. This centromeric region was incorporated to ensure stable maintenance and accurate segregation of the plasmid within *P. murina* cells *in vivo*, mimicking chromosomal behavior and preventing plasmid loss, a common challenge with traditional episomal plasmids in eukaryotic systems in the absence of continuous selective pressure.

**Figure 3.**
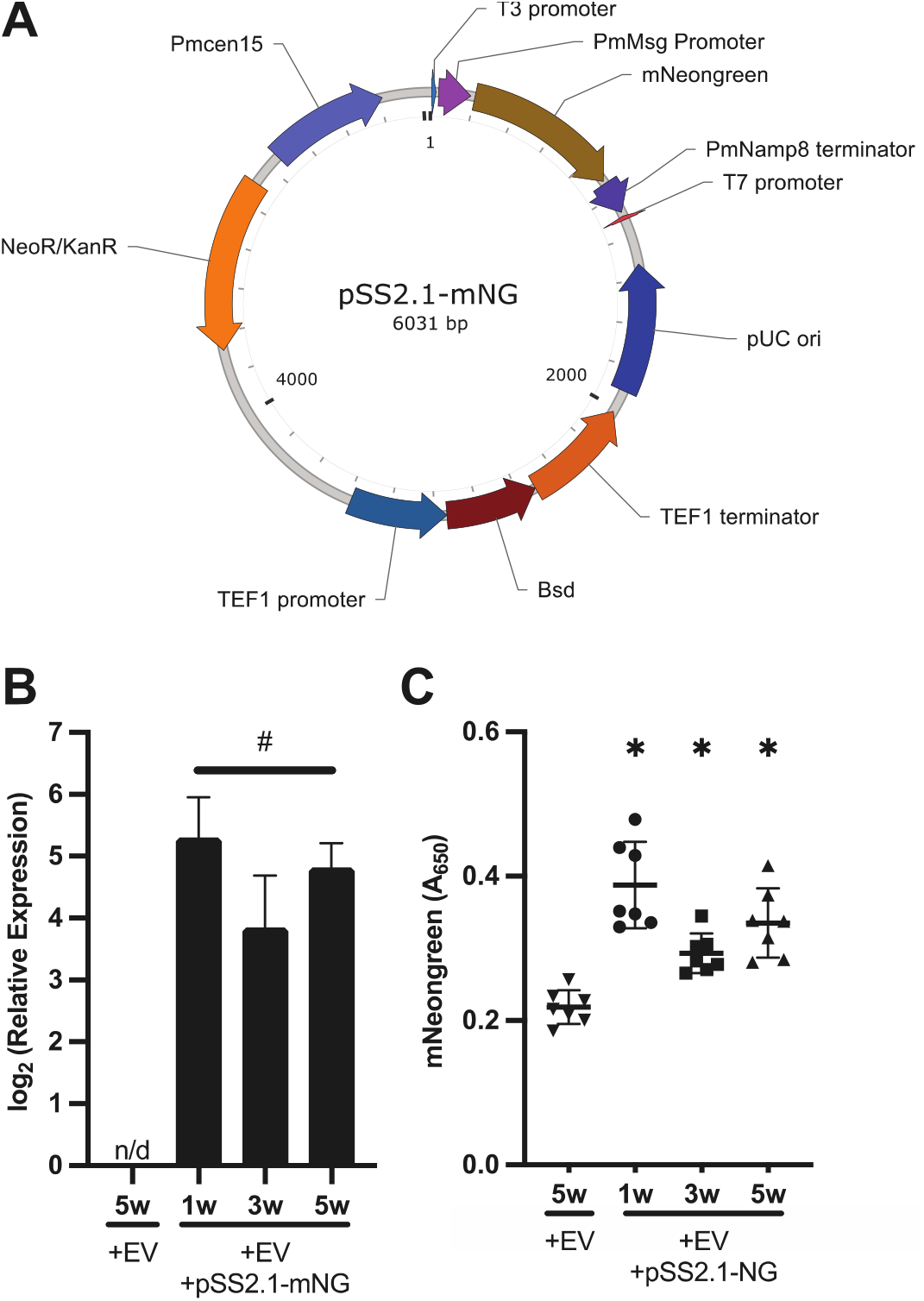
*Pneumocystis murina* expresses mNeonGreen *in vivo* following *in vitro* extracellular vesicle-mediated gene delivery. (A) Plasmid map of pSS2.1-mNG, which includes a second expression cassette encoding blasticidin S deaminase (Bsd) and PmCen15, a truncated centromeric sequence. *P. murina* (1 × 106 organisms) were incubated for 16 hours with extracellular vesicles (2 μg protein equivalent) either lacking cargo or loaded with pSS2.1-mNeonGreen (mNG). Total RNA was extracted, synthesized to cDNA and RT-qPCR was performed. Relative expression was calculated using the 2^-ΔCt^ method, normalized to LSU. (B) Pm displayed sustain expression of *mNeongreen* mRNA expression after 1-, 3-, and 5-weeks post-inoculation (n=3). (C) mNeongreen protein was detected in *P. murina* lysates by ELISA after 1-, 3-, and 5-weeks post-inoculation (n=3). ANOVA followed by Sidak’s multiple comparisons post hoc test; n/d, no detectable expression. ^#^, unable to perform comparison due to the lack of detectable expression. *, p < 0.05 against control.

Following *in vitro* blasticidin selection of *P. murina* transformed with pSS2.1-*mNeongreen*-containing EVs and subsequent infection of mice, *mNeongreen* transcript expression was detected by RT-qPCR and subsequent agarose gel electrophoresis (Figure 3B). While mNeongreen was indetectable by immunofluorescence, likely due to low expression, mNeongreen protein expression was successfully detected from Pm lysates using ELISA (Figure 3C). Both mNeongreen mRNA and protein expression were sustained in lung samples collected at 1 week, 3 weeks, and 5 weeks post-infection. This sustained expression over an extended period clearly indicates successful long-term maintenance and active expression of the pSS2.1 plasmid within the *P. murina* population residing in the host lung environment.

### CRISPR/Cas9 System Enables Targeted Genetic Editing in *P. murina*

Two distinct sets of crRNAs designed to target the *Dhps* gene were validated through *in vitro* cleavage assays. Agarose gel electrophoresis and subsequent densitometry analysis (Figure 4A/B) confirmed efficient cleavage of the *Dhps* target DNA, demonstrating the functionality of the designed crRNAs and the Cas9 enzyme in a cell-free system. This *in vitro* validation was a prerequisite for proceeding with *in vivo* applications.

**Figure 4.**
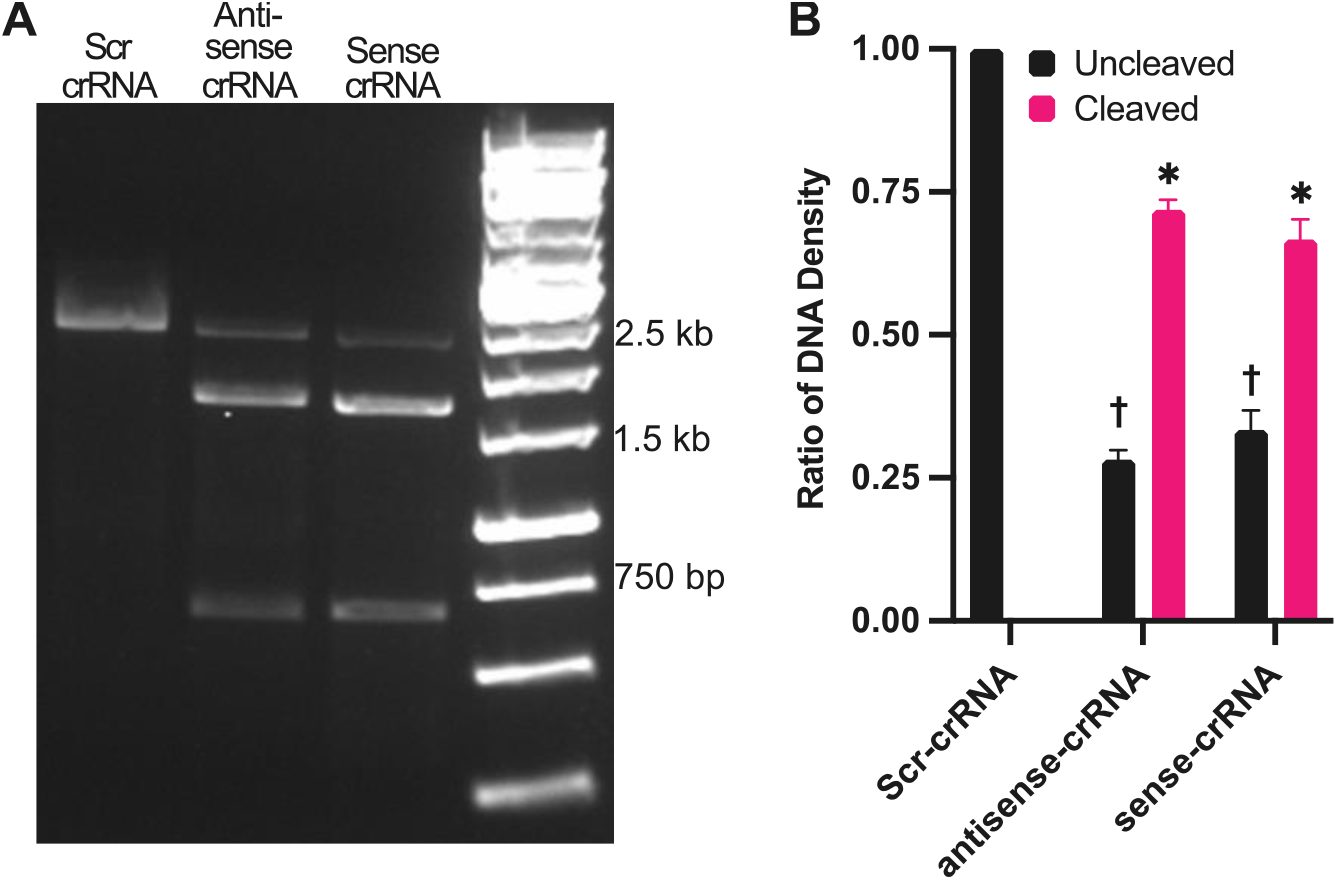
CRISPR RNA (crRNA) sequences mediate efficient cleavage of *Dhps* amplicons in vitro. A full-length Dhps amplicon was generated from *P. murina* genomic DNA. Annealed crRNA and tracrRNA were complexed with *S. pyogenes* Cas9 nuclease to form ribonucleoprotein (RNP) complexes, which were then incubated with the Dhps amplicons for 1 hour. (A) Agarose gel electrophoresis showed cleavage of *Dhps* by both sense and antisense RNPs, yielding two fragments of ∼1560 bp and ∼660 bp, as predicted. Scramble (scr) RNPs did not induce cleavage. (B) Densitometric analysis confirmed significantly increased cleavage by sense and antisense RNPs compared to scr-RNPs (n=3). ANOVA followed by Sidak’s multiple comparisons post hoc test; †, p < 0.05 vs. uncleaved control; *, p < 0.05 vs. cleaved control.

For *in vivo Dhps* editing, we engineered a Cas9 expression cassette encoding a guide RNA targeting the *Dhps* locus, flanked by a hammerhead (HH) ribozyme at the 5′ end and a hepatitis delta virus (HDV) ribozyme at the 3′ end (Figure 5A). This ribozyme-flanked gRNA design was inserted into the CDS region of plasmid pSS2.1 and is based on a previously described self-processing architecture ^24^. In this system, the HH and HDV ribozymes mediate RNA self-cleavage ^25,26^, resulting in the generation of mature Cas9 mRNA and *Dhps*-targeting gRNA. The resulting construct (pSS2.1-Cas9-HH-gRNA^*Dhps*^-HDV) was delivered to *P. murina* via EVs, along with either sense or antisense single-stranded DNA (ssDNA) donor templates encoding the Dhps^ARS^ mutation. Both orientations of the donor were tested to account for potential strand bias during homology-directed repair.

**Figure 5.**
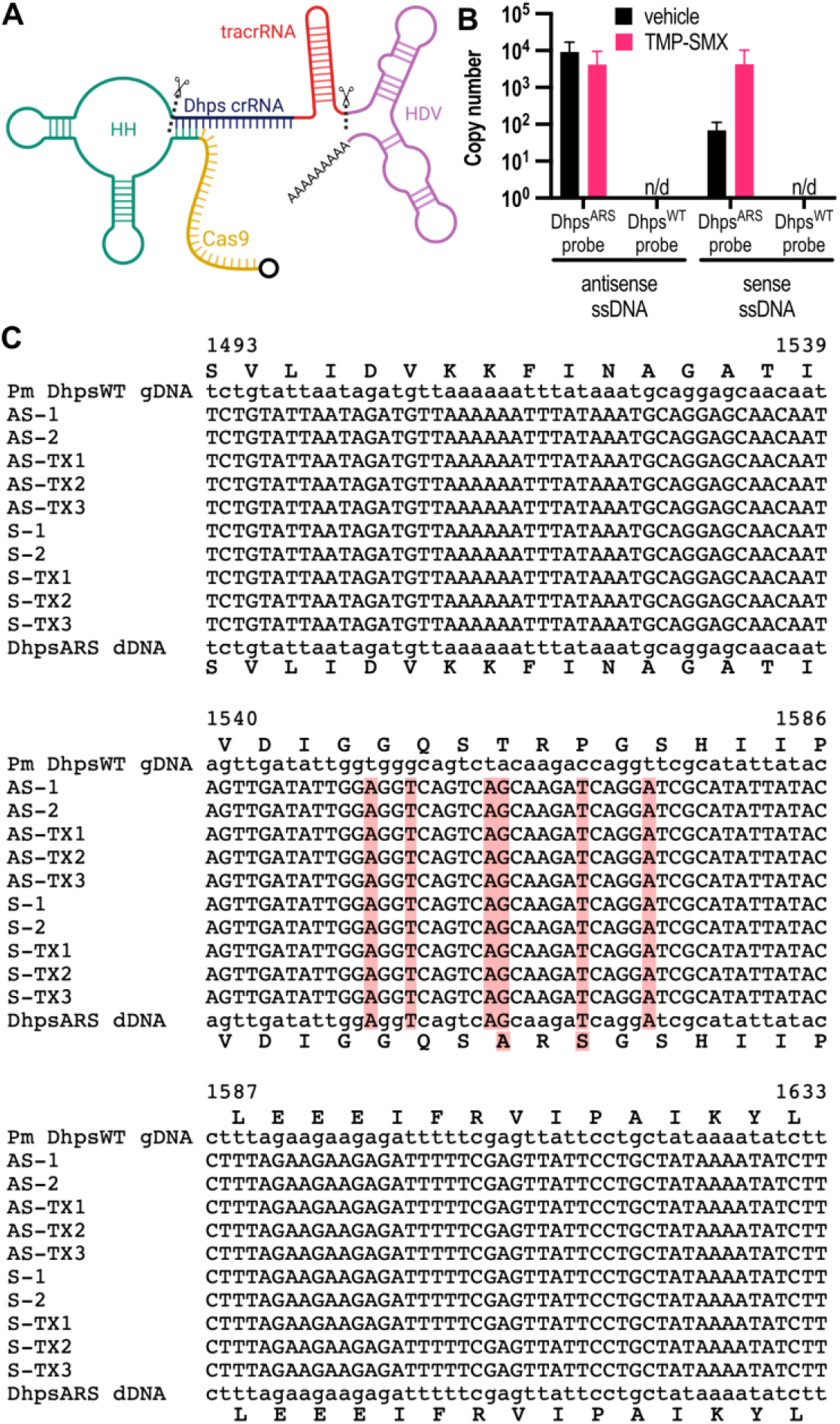
pSS2.1 effectively delivers Cas9 and gRNA for genetic editing of the *Dhps* locus. (A) Schematic of the Cas9-gRNA expression cassette, which encodes Cas9 and a guide RNA flanked by a hammerhead (HH) ribozyme and a hepatitis delta virus (HDV) ribozyme. Created in BioRender. Sayson, S. (2025) https://BioRender.com/dp0twrz. (B) After 5 weeks of infection and 2 weeks of treatment, quantitative PCR detects *Dhps*^*ARS*^ in DNA extracted from *P. murina* in both antisense- and sense-ssDNA-treated groups (n=5). In the same *P. murina* populations, *Dhps*^*WT*^ was not detected. (C) Alignment of the *Dhps* [1493-1633] regions from untreated organisms (Pm DhpsWT gDNA) were compared to *P. murina* organisms treated with EVs containing pSS2.1-Cas9-HH-gRNA^*Dhps*^-HDV and antisense (AS) or sense (S) ssDNA. Donor DNA for Dhps^ARS^ is displayed in bottom row of alignment. Treated Pm organisms all display precise genetic editing to include the associated nucleotide changes for an amino acid change from TRP to ARS, as well as silent point mutations included on the donor dDNA. TX, TMX-SMX groups. Red, change in sequence from DHPS^WT^ isolated from Pm.

Following *in vitro* blasticidin selection to enrich for transformed organisms, mice were infected with the treated *P. murina* organisms. After 5 weeks post-inoculation, the animals were treated with trimethoprim-sulfamethoxazole (TMP-SMX) for two weeks. This treatment regimen applied strong selective pressure, first with blasticidin, for which resistance is encoded on the plasmid, and then with TMP-SMX, where only *Pneumocystis* organisms that had successfully incorporated the Dhps^ARS^ mutation were resistant to SMX exposure ^9-13^.

After 5 weeks of infection and 2 weeks of treatment, quantitative PCR revealed that *Dhps*^*ARS*^ copy number was detectable in both donor-treated groups compared to Dhps^WT^, which was not detected (Figure 5B). These results suggest that delivery of either sense or antisense donor templates promoted maintenance or enrichment of the Dhps locus under selective pressure, consistent with successful *in vivo* editing and expansion of resistant organisms. However, treatment with TMP-SMX did not result in a significant decrease in fungal organisms. However, it’s plausible that blasticidin selective pressure was sufficient in maintaining only *P. murina* organisms that successfully received pSS2.1 and the encoded Bsd resistance.

To confirm this hypothesis, Sanger sequencing was performed on amplified *Dhps* genomic DNA to provide definitive evidence of precise genetic editing. Primers were designed to amplify regions flanking the donor DNA, thereby eliminating the possibility of amplifying residual plasmid or unincorporated donor template. Alignment of the *Dhps* [1493–1633] region from untreated organisms (*P. murina Dhps*^*WT*^) was compared to sequences from *P. murina* organisms treated with EVs containing pSS2.1-Cas9-HH-gRNADhps-HDV and either antisense (AS) or sense (S) single-stranded DNA (ssDNA) donor templates. The alignment (Figure 5C) revealed that 10 out of 10 treated animals harbored fungal organisms with successful homology-directed repair (HDR), displaying precise genomic integration of the *Dhps*^*ARS*^ mutant sequence. In addition to the TRP-to-ARS amino acid substitution, the edited sequences contained silent point mutations engineered into the donor DNA to permit specific qPCR detection and reduce the likelihood of repeated Cas9 RNP binding. Together, these findings provide unambiguous evidence of accurate and durable editing of the *P. murina* genome *in vivo*.

## DISCUSSION

For decades, the inability to culture *Pneumocystis* species *in vitro* has been the principal obstacle to dissecting their biology, metabolism, and pathogenic mechanisms ^5^. This major bottleneck has severely limited the application of modern molecular genetics to this clinically important pathogen. Here, we report an EV-mediated system that bypasses that limitation and establishes the first robust *in vivo* method for genetic manipulation of *Pneumocystis murina*. Because the fungus appears to internalize host EVs naturally, potentially for nutrient acquisition [6], our strategy exploits an existing uptake pathway, which likely underlies the high transformation and transgene-expression efficiencies we observed.

Our study introduces two distinct yet complementary genetic manipulation platforms. The first is a plasmid-based system designed for stable gene overexpression and complementation studies. To achieve sustained gene expression beyond transient delivery, the pSS2.1 plasmid carried a blasticidin-resistance cassette for initial enrichment and a truncated *PmCen15* centromeric region designed to promote episomal maintenance. Consistent with this design, we detected transgene expression for at least five weeks *in vivo*. Although plasmid retention was not directly quantified, consistent mRNA and protein detection across multiple time points supports stable maintenance. These data suggests that inclusion of the centromeric element significantly reduced plasmid loss, thereby permitting longitudinal analyses of gene function and host-pathogen interactions under physiologically relevant conditions. As a plasmid-based expression system, pSS2.1 thus provides a rapid platform for gene overexpression or complementation experiments without permanent genomic modification, enabling researchers to study the effects of increased protein levels or restore gene function in mutant strains. This stable maintenance *in vivo* is a major methodological achievement, opening up new possibilities for conducting prolonged functional studies and investigating drug efficacy over time in a physiologically relevant setting.

Building on that stable plasmid platform, we developed and demonstrated a high-fidelity CRISPR/Cas9 system for precise homology-directed repair (HDR) of the Dhps gene. This system utilized a Cas9 expression cassette with a guide RNA flanked by hammerhead and HDV ribozymes, following the self-processing gRNA architecture described by Gao and Zhao ^24^. The successful *in vitro* cleavage confirmed the functionality of our designed crRNAs and Cas9 enzyme. The subsequent *in vivo* application yielded remarkable results: all ten *P. murina* populations, each derived from independently infected mice, carried the DhpsARS mutation, confirmed by Sanger sequencing. This high efficiency of precise HDR *in vivo* is a groundbreaking achievement for an obligate pathogen. The successful demonstration of HDR in all ten *P. murina* populations is significant, particularly given the general challenges associated with HDR efficiency ^27,28^ and the extreme difficulty of genetically manipulating *Pneumocystis*. This high efficiency of precise HDR is particularly notable, given that *Pneumocystis* appears to primarily rely on homologous recombination (HR) for DNA repair, potentially lacking non-homologous end-joining (NHEJ) pathways ^29^. This indicates that our system successfully directed the organism’s inherent HR machinery for precise genomic modification. This efficiency, coupled with the precision of HDR ^27,28^, means that specific, predetermined genetic changes can now be reliably introduced into *P. murina in vivo*.

The strategic and sequential use of two selective pressures was critical to achieving this success: *in vitro* blasticidin enrichment for initial plasmid uptake, followed by *in vivo* trimethoprim/sulfamethoxazole treatment. This drug pressure amplified the rare precise editing events, providing a reliable route to isolate drug-resistant, site-specific mutants. The ability to precisely introduce the Dhps^ARS^ mutation, a clinically relevant resistance determinant in *P. jirovecii* ^9-13^, directly demonstrates the power of this CRISPR platform. This system complements the plasmid-based approach by enabling stable knock-in, knock-out, or point-mutation studies at native chromosomal loci, allowing for investigations into essential gene functions, protein localization, and the molecular mechanisms of drug resistance.

Together, the stable EV-mediated plasmid transformation for overexpression and the high-fidelity CRISPR editing for precise genome modification furnish complementary tools that finally open *Pneumocystis* research to sophisticated molecular genetics. Investigators can now generate specific drug-resistant strains for *in vivo* drug screening, tag endogenous proteins for localization and interaction studies, or create conditional knockouts of essential genes. These approaches were previously inaccessible for this pathogen. These capabilities are vital for understanding *Pneumocystis* pathogenesis, identifying novel drug targets, and ultimately developing more effective therapies for PjP, especially in the face of emerging drug resistance.

However, the current system has limitations, including the relatively prolonged timeline required. Each genetically edited *P. murina* population necessitates an initial mouse infection lasting approximately 5–6 weeks to yield ∼10^4^ organisms, which then require further propagation in another mouse for an additional 5–6 weeks to achieve larger populations. Additionally, dependence on animal models inherently constrains throughput and scalability compared to traditional *in vitro* cultivation methods. Nonetheless, these limitations are expected to be addressed when a suitable *in vitro* culture system for *Pneumocystis* is developed, which would facilitate selection, scalability, and throughput.

This work delivers a groundbreaking genetic toolkit for *P. murina*, fundamentally transforming the *Pneumocystis* research landscape. These strategies are potentially applicable to other unculturable obligate pathogens facing similar genetic manipulation challenges. By overcoming the long-standing barriers of genetic intractability, this platform significantly advances our ability to investigate *Pneumocystis* biology and pathogenicity at an unprecedented level of molecular detail. By enabling both durable transgene expression and highly efficient HDR *in vivo*, this toolkit paves the way for comprehensive mechanistic studies, precise genetic engineering, and accelerated drug discovery, fundamentally enhancing our capacity to combat *Pneumocystis* pneumonia.

## MATERIALS AND METHODS

### Animals

BALB/c male mice (20-25g; 5-6 weeks) were immunosuppressed for the duration of the study with dexamethasone (4 mg/L) in drinking water, available *ad libitum*. Mice were infected with *P. murina* by intranasal inoculation of 10^6^ organisms. Mice were euthanized humanely and their lungs removed for isolation and quantification of fungal organisms (n=5). To quantify fungal burden, lungs were homogenized in PBS using gentleMACS (Miltenyi Biotec, Auburn, CA, USA), then stained with a modified Diff-Quik staining ^30^ to visualize the nuclei for microscopic enumeration.

These studies were performed in accordance with the Guide for the Care and Use of Laboratory Animals, 8th ed. (National Academies Press, Washington, DC, USA), in AAALAC-accredited laboratories under the supervision of veterinarians. In addition, all procedures were conducted in compliance with the Institutional Animal Care and Use Committee at the Veterans Affairs Medical Center, Cincinnati, OH, USA.

### Extracellular Vesicle (EV) Isolation and Characterization

Mouse lung EVs were isolated and characterized as previously described ^20^. Briefly, BALF was collected from mice using cold PBS (1 mL x 3) and the cellular debris was removed by centrifugation at 3400 x g for 15 minutes. Size exclusion chromatography was performed on BALF using qEV10 columns and the Automatic Fraction Collector (Izon Science, Medford, MA) to isolate EVs from fractions 1-5. EVs were quantified using Micro BCA Protein Assay Kit (Thermo Scientific; Rockford, IL). Quality control measures were performed as previously described ^20^, which included nanoparticle tracking analysis (NTA) for size distribution and concentration, electron microscopy for morphological assessment, and Western blot analysis for common EV markers (e.g., CD9, CD63, TSG101) and absence of cellular contaminants (Data not shown).

### *Pneumocystis murina* maintenance cultures and isolation

*P. murina* organisms were isolated from infected mice and cryopreserved in liquid nitrogen. *P. murina* maintenance cultures were prepared at a density of 1×10^6^ cells/mL using Roswell Park Memorial Institute (RPMI) 1640 medium (Gibco, Grand Island, NY) supplemented with 10% fetal bovine serum (FBS) (Cytiva, Marlborough, MA), 1,000 U/mL penicillin, 1,000 μg/mL streptomycin (Gibco, Grand Island, NY), 1% minimum essential medium (MEM) vitamin solution (Gibco, Grand Island, NY), and 1% MEM non-essential amino acid solution (Gibco, Grand Island, NY) ^31^. Short-term maintenance cultures were distributed in triplicate into 48-well polystyrene plates (Corning Costar, Glendale, AZ) and incubated at 37°C in a 5% CO2 atmosphere.

### Plasmid backbone construction

The initial pSS1 plasmid was designed and constructed from gene fragments. The pSS1 plasmid (Figure 1) was engineered for the transfer and expression of genetic material, encoding either *Dhps*^*ARS*^ or mNeonGreen. These genes were under the transcriptional control of a *P. murina* major surface glycoprotein (*msg*) promoter, which has been previously shown to exhibit strong transcriptional activity in heterologous systems, including *Saccharomyces cerevisiae* ^32,33^. A 151 bp sequence downstream of the 3’ untranslated region of *Namp8* (*PNEG_1673*) was selected and used as a transcriptional terminator.

Subsequent modifications and the generation of various derivative versions, including pSS2.1, were designed in-house and constructed by VectorBuilder. The pSS2.1 plasmid (Figure 4) was further enhanced by the incorporation of a blasticidin resistance gene, blasticidin S deaminase (*Bsd*), for selection. This was under transcriptional control of a *Tef1* (*PNEG_01204*) promoter (444 bp upstream of start codon) and terminator (500 bp downstream of stop codon). Additionally, a truncated PmCen15 centromeric region was added to ensure stable maintenance ^23^.

To promote efficient transgene expression, the coding sequences for *Bsd, mNeonGreen* and *Cas9* were codon-optimized for *P. murina* codon usage using the online codon optimization tool provided by NovoPro (https://www.novoprolabs.com/tools/codon-optimization). The plasmids used in this study are available from Addgene: pSS1-mNeongreen (Addgene plasmid #241014), pSS2.1-Cas9-HH-gRNADhps-HDV (Addgene plasmid #241015), and pSS2.1-mNeongreen (Addgene plasmid #241016).

### EV Transformation and *in vitro* delivery to *P. murina*

Plasmids and siRNA were transformed into purified mouse lung EVs utilizing the Exo-Fect™ Exosome Transfection Kit, following the manufacturer’s protocol (System Biosciences; Palo Alto, CA). Transformation reactions were stopped and EVs were purified using ExoQuick-TC (System Biosciences; Palo Alto, CA) to remove residual nucleic acids, then resuspended in PBS. A 2 μg protein equivalent of loaded EVs or control EVs were introduced to 10^6^ *P. murina* and incubated for a 24-hour period, as described above.

### RNA and DNA Isolation

Total RNA was extracted from *P. murina* cultures utilizing the ZymoResearch Direct-zol RNA Miniprep Kit (Zymo Research, Irvine, California). Genomic DNA was extracted from *P. murina* organisms isolated from lung samples using the ZymoResearch Quick-DNA Miniprep kit (Zymo Research, Irvine, California).

### Reverse transcription quantitative PCR (RT-qPCR) and Quantitative PCR (qPCR)

cDNA was synthesized from isolated RNA using SuperScript™ IV VILO™ Master Mix (Invitrogen, Carlsbad, CA,USA). RT-qPCR was performed on mNeonGreen expression normalized against LSU using PowerUp™ SYBR™ Green Master Mix (Applied Biosystems, Waltham, Massachusetts). RT-qPCR was performed on Dhps^WT^ or Dhps^ARS^ expression normalized against large subunit (LSU) rRNA using TaqMan™ Fast Advanced Master Mix (Applied Biosystems, Waltham, Massachusetts). Real-time PCR assays were run on an Applied Biosciences 7500 Fast PCR system. Relative expression was calculated using the 2^-ΔCt^ method, normalized to the endogenous housekeeping gene *LSU*.

For genomic DNA copy number analysis, qPCR was performed using TaqMan™ Fast Advanced Master Mix. A standard curve was generated using synthetic gene fragments (Azenta Life Sciences, South Plainfield, NJ) of Dhps^WT^ and Dhps^ARS^ to determine absolute copy numbers. Primer, probe, and gene fragment sequences provided in Supplementary Table 1.

### Enzyme-Linked Immunosorbent Assay (ELISA)

The presence of mNeonGreen *in vivo* was detected by ELISA. Lung homogenates were lysed in RIPA buffer (Alfa Aesar; Ward Hill, MA). High-binding 96-well plates (Corning 2592; Kennebunk, ME, USA) were coated overnight with capture antibody, mNeonGreen VHH Recombinant Alpaca Monoclonal Antibody (ChromoTek CTK0203; Planegg-Martinsried, Germany), in 50 mM carbonate buffer, pH 9.4 at 4°C. Lung lysates were diluted 1:10 in PBS and then added to the plate for 1 hour. After antigen capture, wells were washed three times with PBS with 0.1% Tween-20 (PBST), then detected using mNeonGreen Polyclonal Antibody (Proteintech 29523-1-AP; Planegg-Martinsried, Germany) and Goat anti-Rabbit IgG (H+L) Secondary Antibody, HRP (Invitrogen 31460; Waltham, MA). After washing with PBST three times, detection was performed using 1-Step™ TMB ELISA Substrate Solutions (Thermo Scientific; Rockford, IL) and absorbance at 650 nm was read using a Synergy HTX plate reader (BioTek; Winooski, VT).

### CRISPR reagents and *in vitro* cleavage assay

Two distinct sets of crRNA targeting the *P. murina Dhps* gene were designed using the CRISPR tool in Benchling. *P. murina Dhps* was amplified by PCR from gDNA and used for cleavage assays. crRNA was annealed to tracrRNA (IDT; Coralville, IA) at a 1:1 ratio then complexed with recombinant Alt-R *S*.*p*. Cas9 protein (IDT; Coralville, IA). *In vitro* cleavage assays were conducted by incubating cr-/tracr-RNA:Cas9 RNP complexes with *Dhps*^*WT*^ DNA for 60 minutes at 37°C. After the digestion was completed, Proteinase K (1.81 mg; Zymo Research, Irvine, California) was added to stop the reaction. Cleavage products were resolved on a 1% agarose gel in TAE buffer then stained with GelRed Nucleic Acid Gel Stain (Biotium; Fremont, CA). Gels were visualized and analyzed on an iBright CL1500 imaging system (Invitrogen, Carlsbad, CA,USA).

Densitometry analysis was performed employing the iBright software to quantify cleavage efficiency. The specific crRNA sequences and PCR primers are provided in Supplementary Table 1.

### *In vivo* Experimental Design

*P. murina* (1×10^6^) was co-cultured with either untreated EVs or transformed EVs, as described above.

After two hours of incubation, to allow expression of Bsd, blasticidin (100 ug/mL; Gibco; Grand Island, NY) selection was applied *in vitro* for 24 hours. *P. murina* was then washed 3 times in PBS and intranasally inoculation into mice. For pSS2.1-mNeongreen experiments, lung samples were collected at 1 week, 3 weeks, and 5 weeks post-inoculation for subsequent analysis. For CRISPR/Cas9 experiments, mice were allowed to develop infection for 5 weeks, followed by daily treatment with Sulfamethoxazole and Trimethoprim Oral Suspension (125 mg/kg, 12.5 mg/kg; Aurobindo Pharma USA; East Windsor, NJ) three times a week for 2 weeks. Lung samples were collected at the end 2-week treatment period.

### Sanger Sequencing

Genomic DNA was extracted from *P. murina* isolated from mouse lungs and amplified using primers positioned outside the donor DNA regions to ensure amplification of genomic DNA rather than residual donor DNA. The amplified products were submitted to Genewiz for Sanger sequencing. Sequencing primer provided in Supplementary Table 1. Sequence alignments were performed using Clustal Omega to confirm successful homologous recombination ^34^.

### Statistical Analysis

Statistical analysis was performed in GraphPad Prism version 10.4.2 (534) using unpaired t tests or two-way analysis of variance (ANOVA) followed by Sidak’s multiple comparisons post hoc test to control groups. A p-value < 0.05 was considered statistically significant.

## ACKNOWLEDGEMENTS

This work was supported by grants to AGS from the University of Cincinnati, Department of Internal Medicine Senior Pilot Award Program and the NIAID NIH 1R61AI187097.

